# Topographic barriers drive the pronounced genetic subdivision of a range-limited fossorial rodent

**DOI:** 10.1101/2023.04.06.535856

**Authors:** Victoria M. Reuber, Michael V. Westbury, Alba Rey-Iglesia, Addisu Asefa, Nina Farwig, Georg Miehe, Lars Opgenoorth, Radim Sumbera, Luise Wraase, Tilaye Wube, Eline D. Lorenzen, Dana G. Schabo

## Abstract

Due to their limited dispersal ability, fossorial species with predominantly belowground activity usually show increased levels of population subdivision across relatively small spatial scales. This may be exacerbated in harsh mountain ecosystems, where landscape geomorphology limits species’ dispersal ability and leads to small effective population sizes, making species susceptible to environmental change. The giant root-rat (*Tachyoryctes macrocephalus*) is a highly fossorial rodent confined to the afro-alpine ecosystem of the Bale Mountains in Ethiopia. Using mitochondrial and low-coverage nuclear genomes, we investigated 77 giant root-rat individuals sampled from nine localities across its whole ∼1,000 km^2^ range. Our data revealed a distinct division into a northern and southern subpopulation, with no signs of gene flow, and higher nuclear genetic diversity in the south. Landscape genetic analyses of the mitochondrial genomes indicated that population subdivision was driven by steep slopes and elevation differences of up to 500 m across escarpments separating the north and south, potentially reinforced by glaciation of the south during the Late Pleistocene (∼42,000 to 16,000 years ago). Despite the pronounced subdivision observed at the range-wide scale, weak geographic structuring of sampling localities within subpopulations indicated gene flow across distances of at least 16 km, suggesting aboveground dispersal and high mobility for relatively long distances. Our study highlights how topographic barriers can lead to the genetic subdivision of fossorial species, despite their potential to maintain gene flow at the local scale. These factors can reduce genetic variability, which should be considered when developing conservation strategies.

## 1. Introduction

The genetic subdivision and diversity of a species across space are determined by the combined effects of the environment and the ability of a species to disperse (Berthier et al., 2005; Manel et al., 2012; Quaglietta et al., 2013; Ruiz-Gonzalez et al., 2015). Dispersal ability can be limited by topographic barriers such as mountains or steep slopes, and by species-specific abiotic or biotic requirements, such as temperature or food availability, which may prevent the continuous distribution of individuals and reduce gene flow (Boulangeat et al., 2012; Cunningham et al., 2016; Cushman & Lewis, 2010; Sexton et al., 2009). As a consequence, natural selection, genetic drift, and inbreeding in smaller isolated populations may lead to heterogeneous patterns of genetic variability and population subdivision (Wright, 1969). Species with low dispersal ability such as those with low mobility and a burrowing lifestyle, are especially prone to these processes.

Fossorial rodents engineer elaborate underground burrow systems. In many species, activities including searching for mates, reproduction, and foraging, occur below-ground (Nevo, 1999). Therefore, these rodents are often restricted to specific soil types and available food resources (Begall et al., 2007; Nevo, 1999; Reichman, 1975). Apart from these notable constraints in habitat and resource availability, the low mobility of fossorial species leads to small home ranges and limited dispersal (Harestad & Bunnel, 1979; Tucker et al., 2014). As a result, fossorial rodents often have a localised and patchy distribution. Moreover, in the case of solitary species, mature individuals meet mainly during the mating season, which further limits conspecific encounters. Combined, these characteristics lead to small and isolated subpopulations, with low genetic variation and genetic differentiation across relatively small scales and rapid inter-population divergence, as shown for instance in several tuco-tuco species (*Ctenomys sp.,*) and common voles *(Microtus arvalis)* (Mapelli et al., 2012; Mirol et al., 2010; Nevo, 1999; Schweizer et al., 2007).

These genetic and ecological patterns may be exacerbated in harsh environments, such as in mountain ecosystems, where the geomorphology of the landscape and the availability of suitable habitats, limits dispersal opportunities and leads to restricted species distribution ranges and small effective population sizes (Badgley et al., 2017; Brown, 2001; Gaston, 2003; Rahbek, Borregaard, Colwell, et al., 2019). As a result, mountain regions have been recognized as hotspots for genetic differentiation and speciation, contributing disproportionately to terrestrial biodiversity, at least in the tropics (Rahbek, Borregaard, Antonelli, et al., 2019; Rahbek, Borregaard, Colwell, et al., 2019; Sandel et al., 2011). However, species with limited distribution ranges and small population size, such as those found in mountain ecosystems, are at particular risk of extinction (Davies et al., 2009; Gaston, 2003). Small populations tend to exhibit accumulations of deleterious mutations, low intraspecific diversity, or loss of adaptive potential, making them susceptible to environmental change and habitat shifts (Hoffmann et al., 2017; Lande, 1988; Willi et al., 2006). Additionally, upslope habitat shifts are limited for mountain species, especially for those occurring near mountaintops (Parmesan, 2006; Wilson & Gutiérres, 2016). Mountain ecosystems face increasing threats from land use and climate change-induced habitat shifts. Therefore, it is imperative to understand the impact of species-environment interactions on genetic diversity to effectively establish conservation targets, however, thorough understanding is still lacking. Studying species in mountain ecosystems remains a challenge, especially for fossorial rodents, due to the difficulty of accessing remote areas and the inherent challenges in assessing these species in their natural habitats.

Our study addresses this knowledge gap by elucidating how landscape features drive the genetic subdivision and diversity of the giant root-rat (*Tachyoryctes macrocephalus*), a fossorial rodent endemic to the afro-alpine and afro-montane ecosystem of the Bale Mountains in southeast Ethiopia (Figure 1). The species has a limited distribution range of ∼1,000 km^2^ across the Bale Mountains massif and is found between 3,000 and 4,150 m above sea level (a.s.l.) (Sillero-Zubiri et al., 1995; Yalden & Largen, 1992). Giant root-rats have specific habitat requirements, occurring in grasslands in areas with good soil depth, especially along wetland shores and flooded valleys (Sillero-Zubiri et al., 1995; Šklíba et al., 2017). Grassland in river valleys that spread through shrubs and forest zones into lower elevations, allow the species to expand down to about 3,000 m a.s.l. (Yalden, 1985). Their relatively small home ranges (about 100 m^2^) can shift throughout the year depending on food availability (Šklíba et al., 2020). Giant root-rats are significant ecosystem engineers creating large underground burrow systems, in which they live solitarily. Through their combined effect of soil perturbation and herbivory, they alter nutrient availability, soil texture and moisture, and create their own habitat and that for other plant and animal species (Asefa et al., 2022; Miehe & Miehe, 1994; Šklíba et al., 2017; Yalden, 1985). By using below-ground burrows, the species circumvents the harsh environmental conditions of the mountain ecosystem, which include strong winds and temperatures below 0 C°, and limits the risk of being preyed upon by its main predator the Ethiopian wolf (*Canis simensis*) (Sillero-Zubiri & Gottelli, 1995; Šumbera et al., 2020; Vlasatá et al., 2017; Yalden, 1985). Taken together, the species’ key role as ecosystem engineer combined with its limited range in a changing mountain ecosystem, makes it an ideal model organism for investigating the connection between genetic patterns and landscape features, so as to preserve mountain biodiversity and ensure ecosystem functioning.

**Figure 1:**
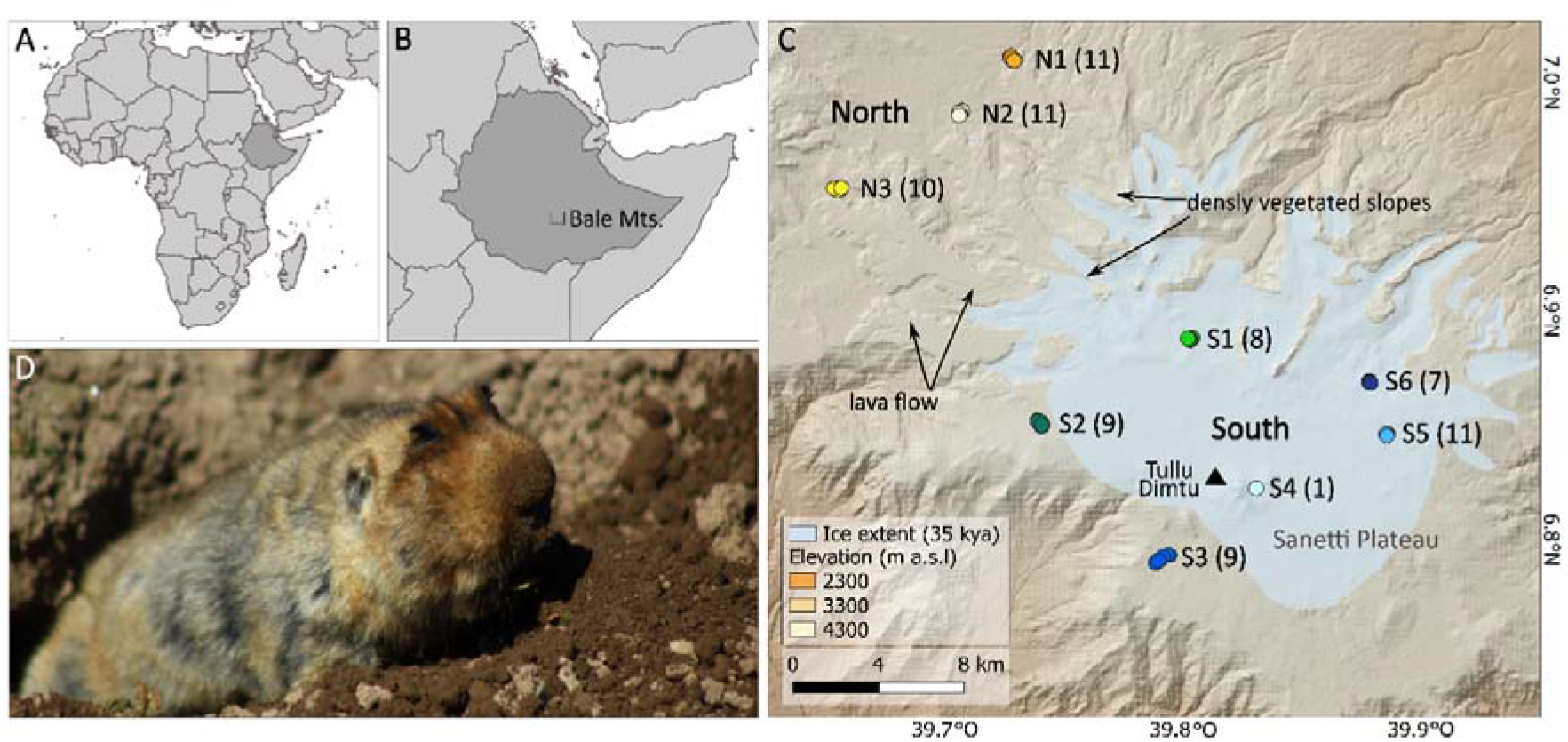
Sampling localities of giant root-rats within Bale Mountains National Park, Ethiopia. A) Map of Africa showing Ethiopia in dark grey; B) Map of Ethiopia indicating location of Bale Mountains National Park, C) Map of the nine sampling localities from two distinct geographic regions. The north (∼3,500 m above sea level [m a.s.l]), and south (∼3,800-4,000 m a.s.l, Sanetti Plateau) are separated by steep slopes covered with dense Erica thickets and congealed lava flows. Sample size of each locality is indicated in brackets. Tullu Dimtu is the highest peak in the Bale Mountain National Park at 4,377 m a.s.l., and is indicated with a filled triangle. Region with light blue shading indicates the glacial extent within Bale Mountains National Park ∼35±7.1 thousand years ago (kya) (Groos et al., 2021; Ossendorf et al., 2019); D) Burrowing giant root-rat, photography by V. Reuber.

In the present study, we analysed the spatial genetic subdivision and diversity in the giant root-rat across its distribution range. To achieve this, we analysed both mitochondrial genomes (mitogenomes) and nuclear genomes, and further utilized mitogenomes to investigate the relationship between genetic differentiation and landscape features. We generated complete mitogenomes and low-coverage nuclear genomes from 77 individuals collected across nine sampling localities in the Bale Mountains (Figure 1). We applied two different landscape genetic approaches to evaluate how mitochondrial gene flow of the species is impacted by geographic distance, by vegetation and soil moisture (used as proxies for food and soil availability), and by slope and elevation (used as proxies for topographic barriers). Due to the predominantly below-ground activity and patchy distribution of the giant root-rat, we hypothesise strong genetic subdivision across small spatial scales. Owing to the pronounced heterogeneity of the environment in the Bale Mountains, we hypothesise that genetic structuring is driven by habitat availability, and by topographic structures across the species’ range.

## 2. Materials and method

### 2.1 Study area

The Bale Mountains in southeast Ethiopia (6°29’N – 7°10’N and 39°28’E – 39°57’) represent Africa’s largest afro-alpine ecosystem, comprising ∼8 % of the continent’s area above 3,000 metres above sea level (m a.s.l) (Groos et al., 2021, Figure 1A-C). In order to protect the unique afro-montane and afro-alpine ecosystem of the Bale Mountains, the area above ∼3,200 m a.s.l. became a national park in 1970. The Bale Mountains are characterised by two rainy seasons and one dry season per year, with short rains from March to June, long rains from July to October, and a dry season from November to February. The vegetation of the Bale Mountains shows an elevational zonation from moist montane forest (∼1,500 - 3,500 m a.s.l.) over ericaceous shrubland and dwarf forest (∼3,500 - 4,000 m a.s.l.) to afro-alpine vegetation with open grassland and *Erica* outposts (above 4,000 m a.s.l.). Dwarf-scrub vegetation, such as *Helichrysum* associated with *Lobelia*, is the main plant formation in the afro-alpine vegetation but does not cover the whole area, leaving open spaces for herbaceous plants like *Senecio*, *Alchemilla* or *Salvia* (Miehe & Miehe, 1994; Tallents & Macdonald, 2011)

Characteristic of the Bale Mountains National Park is the afro-alpine Sanetti Plateau, which spans elevations from approximately 3,800 m a.s.l, to 4,377 m a.s.l. at the peak of the mountain Tullu Dimtu (Figure 1C). Large parts of the plateau were glaciated during the Late Pleistocene, between 42,000 to 16,000 years ago (Groos et al., 2021). The plateau is bounded by several outlet valleys in the north and the east with slopes that are covered by dense, shrubby *Erica* vegetation and by congealed lava flows at its northwestern margins. These topographic structures distinguish the plateau from the northern region of the national park, which is ∼300 - 500 m lower in elevation and comprises broad valleys and plains with afro-alpine vegetation (Miehe & Miehe, 1994). In comparison to the plateau, the north has higher moisture availability and milder temperatures.

### 2.2 Sampling

We collected tissue samples from 77 live giant root-rat individuals at nine localities across the Bale Mountains National Park, covering the distribution range of the species (Figure 1C, supporting information Table S1). The sampling localities were distributed across the two topographically distinct regions in the national park, in the north (localities N1-N3) and in the south (localities S1-S6). The southern localities are scattered across the centre and south of the Sanetti Plateau. Localities sampled in the north of the plateau lie at a lower elevation (∼3,500 m a.s.l) than localities sampled in the south (∼3800 - 4000 m a.s.l.). Sampling localities between regions were separated by 15.3 to 28.3 km, and localities within regions were separated by 2.6 km to 16.0 km.

We captured 7-9 giant root-rat individuals per locality (except locality S4 with n=1). The samples were collected in January and February in two consecutive years (2020, 2021) under the permit of the Ethiopian Wildlife Conservation Authority. Individuals were caught with snare traps that were monitored by the capture team at all times to guarantee no harm to the animals. A ∼0.5 cm^2^ piece of skin from the hind leg was cut with sterilised scissors and stored in 96% ethanol or DNAgard^®^ for blood and tissue (Biomatrica, Inc.) for genomic analyses. After sterilising the wound, the animals were immediately released back into their burrow systems.

### 2.3 Laboratory analyses

We extracted DNA from the tissue samples using the Qiagen DNeasy^®^ Blood and Tissue Kit following the manufacturer’s protocol (Qiagen Ltd.). 60 of the samples were processed in-house in the modern DNA labs at Globe Institute, University of Copenhagen. The DNA concentration of the extracts was measured using QubitTM dsDNA HS (Invitrogen). After quantification, we diluted the extracts to a concentration of 6 ng/µl in a total volume of 50 µl. DNA was sheared to ∼400 base pair (bp) fragment lengths using the Covaris M220 ultrasonicator. We built DNA fragments into an Illumina library following the protocol from Carøe et al., (2018) and double-indexed them using AmpliTaq Gold Polymerase (ThermoFisher) during the indexing PCR step. Index PCR reactions were performed in 100 µl, using 1x PCR buffer, 2.5 mM of MgCl (25 mM), 0.2 mM of dNTPs (25 mM), 0.2 μM of index primer mix (10 µM), and 0.1 U/ µl of polymerase (5 U/µl). PCR cycling conditions were 95 °C for 10 min; 10 -18 cycles of 95 °C for 30 s, 50 °C for 30 s and 72 °C for 1 min followed by 72 °C for 5 min. The number of cycles for the index PCR was determined from qPCR analysis. Post-amplification, libraries were purified using SPRI beads as in Carøe et al., (2018). The purified indexed libraries were quantified on a QubitTM dsDNA HS (Invitrogen), and quality-checked on either an Agilent 2100 Bioanalyser or an Agilent Fragment AnalyzerTM. Libraries were pooled equimolarly and sequenced on an Illumina NovaSeq 6000 using paired-end (PE) 150 bp technology (Novogene Europe, http://en.novogene.com).

For the remaining 17 samples, DNA was extracted and processed to libraries by Novogene and sequenced on a NovaSeq 6000, using paired end 150 bp technology.

### 2.4 Data generation of mitochondrial and nuclear DNA

We trimmed adapters and removed reads shorter than 30 bp for each individual using skewer v0.2.2. (Jiang et al. 2014). We merged overlapping paired-end reads using FLASHv1.2v11 (Magoč & Salzberg, 2011) with default parameters. We mapped both merged and unmerged reads to the hoary bamboo rat (*Rhizomys pruinosus*) nuclear genome (Genbank accession: VZQC00000000.1; Guo et al., 2021) which is the nearest relative of the giant root-rat with an available genome, combined with the giant root-rat mitogenome (Genbank accession: MW751806; Reuber et al., 2021). We used BWA v0.7.15 (Li & Durbin, 2009) utilising the mem algorithm and default parameters. We parsed the alignment files, and removed duplicates and reads of mapping quality score <30 using SAMtools v1.6 (Li et al., 2009). We built consensus mitogenomes from each individual using a majority rules approach (- doFasta 2) in ANGSD v0.921 (Korneliussen et al., 2014) only considering bases with a base quality score greater than 30 (-minq 30), reads with a mapping quality score greater than 30 (-minmapq 30), and sites with at least 10x coverage (-minInddepth 10). The mitogenomes are available under GenBank accessions OQ207545 - OQ207620.

### 2.5. Genetic subdivision

#### 2.5.1 Mitochondrial DNA analysis

##### Haplotype network

The mitogenomes were aligned with Mafft v.7.392 (Katoh & Standley, 2013). We constructed a median-joining haplotype network to investigate the relationships among the 77 mitogenomes using the software PopArt v.1.7 (Leigh & Bryant, 2015).

##### Phylogenetic analysis

We constructed a Bayesian phylogeny with the 48 mitogenome haplotypes identified in the network analyses, using MrBayes v.3.2.7a (Ronquist & Huelsenbeck, 2003). We used the GTR + I + G model of evolution, which was defined as the best model with PartitionFinder v. 2.1.1 (Lanfear et al., 2017) prior to the analysis. The MCMC algorithm was run twice with four chains of 10 million generations, sampled every 1,000 generations and with a 10 % burn-in. The trees were combined following the majority-rule consensus approach, to assess the posterior probability of each clade. The resulting tree was visualised in FigTree v.1.4.4 (http://tree.bio.ed.ac.uk/software/figtree/; supporting information Figure S1).

##### Fixation statistics and AMOVA

We calculated the pairwise differentiation between eight of the nine sampled localities (omitting S4 as n=1) with the F_ST_ -estimator of the software Arlequin v. 3.5.2.2 with 10,000 permutations (Excoffier & Lischer, 2010); *p*-values of F_st_ estimates were adjusted using the Bonferroni correction (Rice, 1989), controlling for a false-positive discovery rate (R Core Team, 2021). Additionally, we conducted a hierarchical analysis of molecular variance (AMOVA), also in Arlequin, with localities grouped into their provenance in the regions north and south (Figure 1C).

#### 2.5.2 Nuclear DNA analysis

We investigated population subdivision of the nuclear data using principal component analysis (PCA) and admixture proportion analysis. We generated genotype likelihoods in ANGSD (Korneliussen et al., 2014) for all individuals using the following filters and parameters: call genotype likelihoods using the GATK algorithm (-GL 2), output a beagle file (-doGlf 2), only include reads with mapping and base qualities greater than 30 (-minmapQ 30 and -minQ 30), only include reads that map to one location uniquely (-uniqueonly 1), a minimum minor allele frequency of 0.05 (-minmaf 0.05), only call a SNP if the p-value is greater than 1e^-6^ (-SNP_pval 1e-6), infer major and minor alleles from genotype likelihoods (-doMajorMinor 1), only include sites if at least 40 individuals are covered (-MinInd 40), remove scaffolds shorter than 100 kb (-rf), and remove secondary alignments (-remove_bads 1). To compute the PCA, we constructed a covariance matrix from the genotype likelihoods using PCAngsd v0.98 (Meisner & Albrechtsen, 2018). Admixture proportions were calculated using the same genotype likelihoods with NGSadmix (Skotte et al., 2013). We ran NGSadmix specifying K=2 and K=3. To evaluate the reliability of the NGSadmix results, we ran each *K* up to 100 times independently. If we retrieved consistent log-likelihoods from at least two independent runs, the corresponding K was considered reliable.

To estimate levels of differentiation among localities, we computed F_ST_ from a consensus haploid call file created using ANGSD (-dohaplocall 2) and the same filtering parameters as the PCA and admixture proportions above. We calculated the F_ST_ using an available python script https://github.com/simonhmartin/genomics_general/blob/master/popgenWindows.py) and specifying a window size of 1Mb, and a minimum number of sites per window as 1,000 bp.

##### Gene flow

The mitogenomes of two individuals (WM07 from locality S1 and GG01 from locality S6) grouped with individuals from the north in the haplotype network and phylogeny. We therefore used D-statistics (also known as ABBA/BABA, Durand et al., 2011) to test whether the results were driven by ancient gene flow between regions north and south, or by incomplete lineage sorting. We tested several topologies [[H1, H2], H3], with branch H1 being one of the two putative introgressed individuals, and branches H2 and H3 being individuals from one region north or south, or one from each region. A negative D-score illustrates a closer relationship between H1 and H3 than H2 and H3, while a positive D-score indicates that branches H2 and H3 are more closely related than H1 and H2. This setup can also be used to uncover population subdivision, as the incorrect input topologies would lead to elevated D-scores due to more recent common ancestry, as opposed to gene flow (Westbury et al., 2018).

We performed the D-statistic tests using a random base-call approach in ANGSD (-doAbbababa 1). We implemented the same filtering approach as for the above analyses but only included scaffolds >1 megabase (Mb) in size, a block size of 1 Mb (-blocksize 1000000), and the hoary bamboo rat (*Rhizomys pruinosus*, GenBank accession VZQC00000000.1, Guo et al., 2021) as the ancestral state/outgroup (-anc). To assess the significance of our results we used a block jackknife test with the script jackKnife.R which is available with the ANGSD tool suite.

### 2.6 Diversity

Based on the mitochondrial genomes, we calculated nucleotide diversity (π) per region, and separately for eight of the nine localities using DnaSp v.6 (omitting S4 as n=1) (Rozas et al., 2003). We tested differences in nucleotide diversity between the two geographic regions, and among localities using genetic_diversity_diffs v. 1.0.3 (Alexander, 2017).

The python script used to calculate nuclear F_ST_ above simultaneously computes nucleotide diversity per region and per locality. To test for significant differences in levels of nucleotide diversity between regions and between localities, we used a Welch-test (unpaired t-test), accounting for unequal variance.

### 2.7 Landscape genetic analysis

We applied two landscape genetic approaches to investigate the effects of landscape features on the observed genetic differentiation between localities based on F_ST_ estimates of the mitogenomes. We exclusively used mitogenomes due to their higher mutation rates and lack of recombination compared to nuclear genomes, which result in faster responses to environmental changes and increased resolution (Avise, 2000; Birky et al., 1983). The limited number of sampling localities prevented us from analysing the north and south regions separately.

We selected four environmental variables (vegetation, soil moisture, slope, elevation), which were based on satellite-based remote sensing data, as predictors for genetic differentiation of the giant root-rat. For vegetation and soil moisture, we used observations from satellite Sentinel-2, captured on an almost cloud-free day on December 15, 2017, and derived from the USGS Earth Explorer repository. We used the red, green, blue, red and near infrared bands of from the Sentinel-2 observations and with those computed raster layers of the Normalised Differentiation Vegetation Index (vegetation index) and Land-Surface Water Index (soil moisture index, for details see Wraase et al., 2022) using the Rtoolbox, as proxies for food and soil availability for giant root-rats (Sillero-Zubiri et al., 1995; Šklíba et al., 2017; Yaba et al., 2011; Yalden & Largen, 1992). To determine whether topographic structures act as barriers for burrowing giant root-rats, we included the variables slope and elevation in our analysis. Raster layers for slope and elevation were obtained from a Shuttle-Radar-Topography-Mission digital elevation model from the USGS Earth explorer (www.earthexplorer.usgs.gov). The generated raster layers of all four environmental variables had a 30 x 30 m resolution and were cropped on extent 567910.0, 605990.0, 738620.0, 778750.0. Our analyses were conducted in R environment version 4.2.1 (R Core Team, 2021).

#### Partial Mantel tests

Using partial Mantel tests, we analysed if the genetic differentiation between eight of the nine sampling localities (omitting S4 as n=1) was correlated with geographic distance, vegetation, soil moisture, slope and elevation (see above). Therefore, we constructed distance matrices. The genetic distance matrix was generated by linearizing the pairwise genetic differentiation estimates between localities, i.e the F_ST_ -estimates ([F_ST_ /(1- F_ST_)]; Rousset, 1997). The geographic distance matrix was calculated in Euclidean distances and log-transformed to linearize the relationship with genetic distance. For each environmental variable, we extracted their values from the computed raster layers at the coordinates of the sampling localities and therewith generated the environmental distance matrices. We then applied a pairwise reciprocal causal modelling approach. Reciprocal causal modelling compares partial Mantel tests of a focal environmental model, removing the influence of a competing, alternative model (Cushman et al., 2006; Cushman & Landguth, 2010). In this approach, the correlation between one environmental distance matrix and genetic distance is controlled by a second matrix (e.g. focal model: genetic distance∼geographic distance|elevation distance) and in a next step, both environmental distance matrices are interchanged (e.g. alternative model: genetic distance∼elevation distance|geographic distance). In that way, we were able to account for high correlation among matrices (Cushman et al., 2006; Cushman & Landguth, 2010). To assess which of the two models explains genetic distance better, the relative support of the focal and alternative model was calculated by estimating the difference between the correlation values of the two models. If the difference in correlation factors was positive, we assumed that the focal hypothesis was correct. The partial Mantel tests were performed with 9,999 permutations in the vegan R package v.2.6-4. (Oksanen et al., 2020).

#### Raster layer optimization framework to generate resistance surfaces

We used a raster layer optimization framework developed by Peterman et al. (Peterman, 2018; Peterman et al., 2014), to further identify landscape features that explain mitochondrial genetic differentiation, using the R package ResistanceGA (Peterman, 2018). In this framework, the raster layers of the environmental variables (see above) were transformed into resistance surfaces, with the ResistanceGA package utilising a genetic algorithm from the GA R package (Scrucca, 2013). A resistance surface is a spatial layer that assigns values to each grid in the raster layer of the selected environmental variable. Those values are used to estimate the cost of dispersal and mirror to what extent the selected variable hinders or facilitates the connectivity of a species between two localities (pairwise resistance distances). Thereby, there are no *a priori* assumptions about the relationship between the environmental variable and the species’ dispersal characteristics. The genetic algorithm in the optimization framework is used to maximise the relationship between the resistance distances of each raster layer, and the pairwise genetic differentiation (F_ST_) between localities. The process of generating resistance surfaces is repeated, and in every iteration, the resistance distances are fitted against the genetic distances in a mixed effect model, until the objective function, the AIC (Akaike’s information criterion; Akaike, 1974) of the mixed effect model does not improve further. The mixed-effect models are conducted using a maximum likelihood population effect parameterization to account for the non-independence of the predictor variables and to account for spatial autocorrelation (Clarke et al., 2002; Peterman et al., 2014; Shirk et al., 2018). This iterative process works towards identifying the best-fit landscape resistance surface.

In our optimization framework, we used a single surface optimization approach, where the resistance surfaces of the raster layers of each selected environmental variable (i.e. vegetation index, soil moisture index, slope and elevation) were optimised individually, using eight transformation functions (Monomolecular and Ricker functions) and the default parameters (Peterman, 2018). In this step, the pairwise resistance distance between localities was estimated by assuming that individuals can use several paths to disperse. Resistance distances were generated with the *costDist* function implemented in the ResistanceGA package and movements between localities were allowed in eight directions during resistance distance calculation. Because Euclidean distance is incorporated in the resistance distances, it was not included as an additional variable. In the optimization process, the mixed-effect models were calculated, fitting the pairwise genetic differentiation as a response against the resistance distances as single fixed effects, using the AIC for model evaluation and including sampling localities as a random effect to account for spatial autocorrelation. We did two independent optimization runs to confirm convergence across runs. The run containing the mixed-model with the greatest log-likelihood value is presented in the results section (Table 1). After the optimization, we used bootstrap model selection with 75% of the samples and 10,000 iterations. The bootstrap model selection refits the mixed-effect models and calculates fit statistics for each model, showing the average AIC and percentage each resistance surface has been selected as a top rank model across all bootstrap iterations (Peterman, 2018).

**Table 1:** Model selection results for linear mixed-effect models, testing the effect of resistance distances (dispersal costs based on environmental condition) on levels of mitochondrial genetic differentiation (F_ST_) in the giant root-rat. The parameters were calculated based on 10,000 bootstrap iterations using a random resampling of 75% of the sampled populations. Frequency top-model percentage, higher average weight and log-likelihood (LL), and lower average rank indicate the best supported models.

| Resistance distance matrix | Average AIC | Average $R^2_m$ | Average LL | Average weight | Average rank | Frequency top model (%) |
| --- | --- | --- | --- | --- | --- | --- |
| Slope | -16.61 | 0.92 | 12.31 | 4.8E-05 | 1.36 | 70.87 |
| Elevation | -10.29 | 0.86 | 9.15 | 0.004 | 2.88 | 29.13 |
| Vegetation | -7.16 | 0.78 | 7.58 | 4.0E-07 | 3.19 | 0 |
| Soil moisture | -6.26 | 0.78 | 7.13 | 1.2E-06 | 3.57 | 0 |
| Distance | -4.20 | 0.65 | 4.10 | 0.990 | 4.00 | 0 |

## 3. Results

We generated complete mitogenomes from all 77 giant root-rat individuals, with a depth of coverage ranging between 24.94x and 383.22x, and a length of 16,646 bp. For the nuclear genomes, we obtained coverages ranging from 0.12x to 0.77x (supporting information Table S2).

### 3.1 Genomic analysis

#### Haplotype network and phylogeny

Our 77 sampled giant root-rat individuals comprised 48 haplotypes, with no haplotypes shared among localities (Figure 2A). The haplotype network and phylogeny of the mitogenomes revealed two geographically separated and well-supported groups; one group comprising all samples collected in the north, with the inclusion of individuals WM07 and GG01 from the south, and one group comprising all other samples from the south (supporting information Figure S1). In the network, we identified 574 segregating sites among individuals, and the northern and southern haplogroups were separated by 334 segregating sites (Figure 2A). The north contained 21 haplotypes (32 individuals) and the south contained 27 haplotypes (45 individuals). Within the north, we identified two distinct, well-supported genetic clades, N_A_ and N_B_ (supporting information Figure S1). Clade N_A_ comprised six individuals (five haplotypes) from localities N1-N3. Individual GG01 from locality S6 was basal to the clade. Clade N_B_ comprised the remaining individuals from the north with individual WM07 from the south at basal position. We did not identify any spatial genetic structuring in the south.

**Figure 2:**
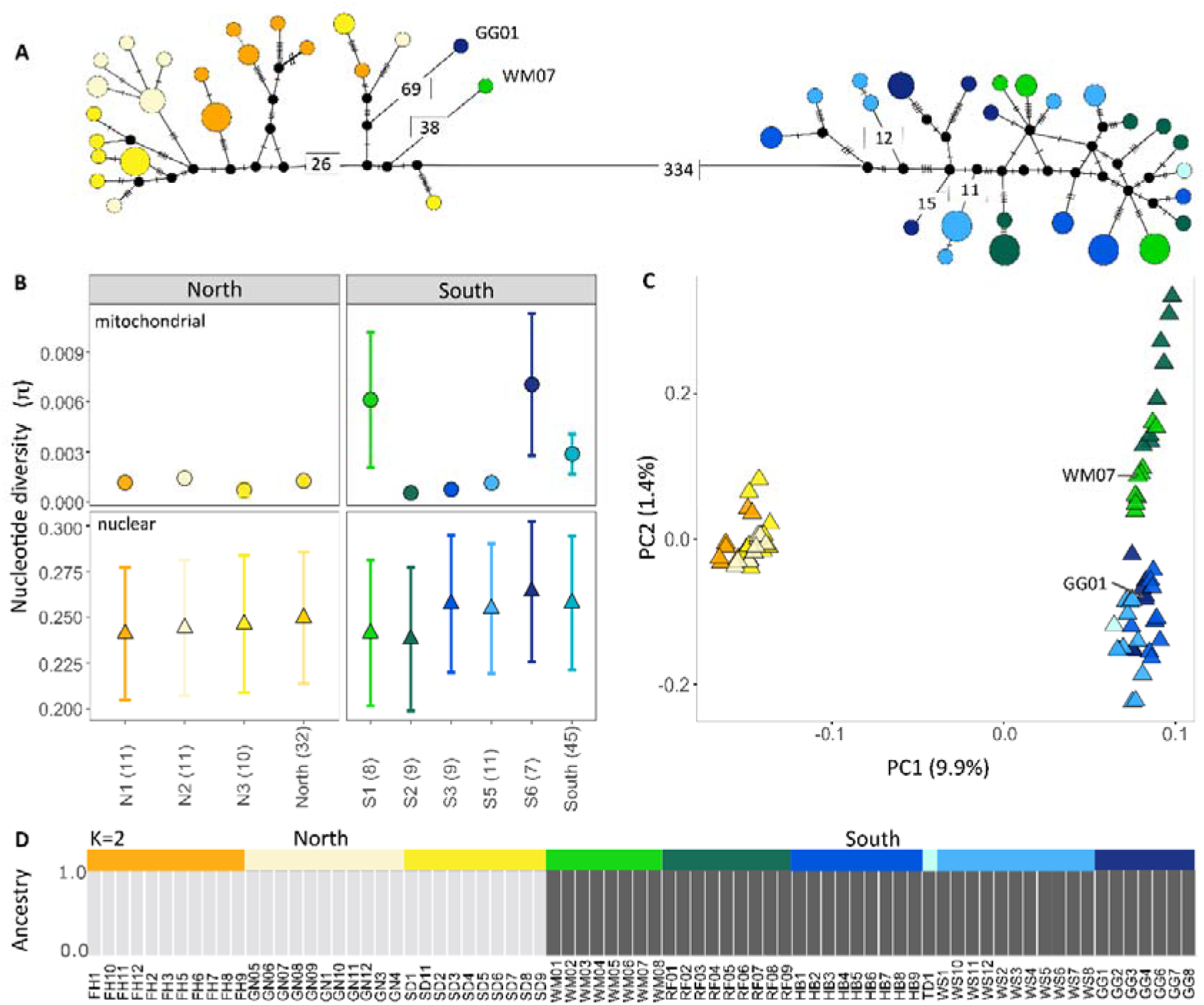
Patterns of genetic diversity and subdivision of giant root-rats across their range. A) Network of the 48 mitochondrial haplotypes present among the 77 sampled individuals. Each circle represents a haplotype, and the relative size of each circle represents haplotype frequency. Numbers on branches show the number of segregating sites between haplotypes for >10. Black dots indicate intermediate haplotypes not present in the data. Distances between haplotypes are not to scale. B) Diversity levels within the sampled localities and subpopulations based on mitogenomes (circles) and nuclear data (triangles). Sample sizes are shown in brackets. C) PCA based on 77 low-coverage nuclear genomes. The percentage of the diversity of each principal component is shown in brackets. D) Ancestry proportions based on the nuclear data for K=2, with each vertical bar representing an individual.

The AMOVA yielded a high level of between-region variation when the eight localities (omitting S4 with n=1) were grouped into north and south (89.50 %, p < 0.01). Within each region, the variation was higher within localities (9.27 %) than among localities (1.24 %).

#### Principal component analysis and admixture proportions

We identified two main groups in the nuclear data, in agreement with the mitochondrial findings (Figure 2). In the PCA, individuals from the north separated from the individuals from the south, with almost 10% of the variation explained on the first principal component. The southern group showed a slight separation on the second component, with the more central localities S1 and S2 segregating from the localities further southeast, with 1.4 % variation explained (Figure 2C). The division of the data into north and south was also evident in the admixture analysis of K = 2 (Figure 2D). The admixture analysis did not converge with K = 3, suggesting K = 3 did not reliably fit the data.

#### Gene flow

To investigate the origin of the mitochondrial lineages, present in individuals WW07 and GG01, which were more closely related to the northern haplogroup than the south (Figure 2A), we tested for ancient gene flow using the nuclear data. Using the topology [[WM07, south], north], we found most comparisons to have a D-score around 0 and a Z-score<|3|, indicating that WM07 and all individuals from the south were equally related to the northern individuals (Figure 3). We found positive D-scores (Z-score>3) when using the topology [[WM07, north], north], demonstrating a closer relationship between individuals from the north with each other than with WM07, which agrees with their more recent common ancestry and the basal position of WM07 in the phylogenetic tree (supporting information Figure S1). We found qualitatively the same results when we investigated the relationship of GG01 (sampled in locality S6) with individuals from the north and south (Figure 3). Hence our analysis did not support that the mitochondrial lineages in WM07 and GG01 were the result of recent gene flow between north and south.

**Figure 3:**
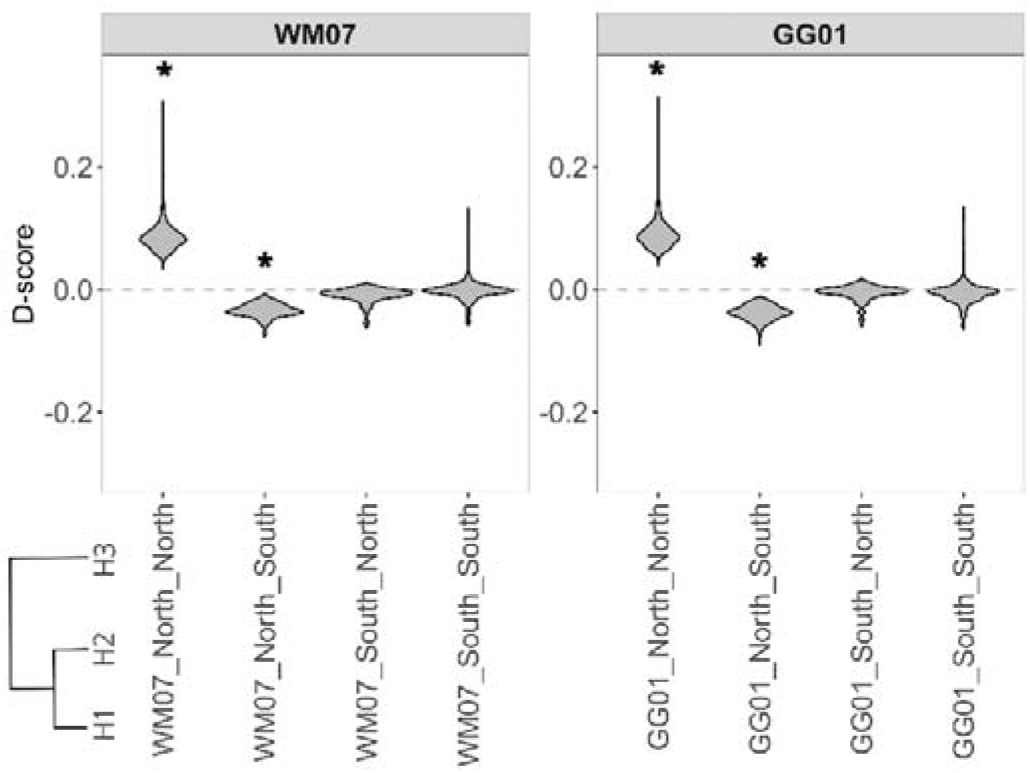
Analysis of signals of gene flow between individuals WM07 from locality S1 and GG01 from locality S6 and their source group in the south, using D-statistics. Negative D-scores suggest gene flow or recent common ancestry between H1 and H3 relative to H2 and H3, while positive D-scores suggest gene flow or recent common ancestry between H2 and H2 relative to H1 and H3. Statistical significance is indicated by asterisks (*) next to scores, when |Z| greater than 3 (supporting information Figure S2), determined by a one-sample Wilcoxon signed rank test.

#### Genetic differentiation among localities

Investigating pairwise genetic differentiation between localities, we found that F_ST_ - estimates were higher between regions than within regions, at both the mitogenome and nuclear level (Figure 4). For the mitogenomes, pairwise differences between localities of different regions ranged from 0.82 to 0.97 and were much higher than within regions, where values ranged from 0.09 to 0.29.

**Figure 4:**
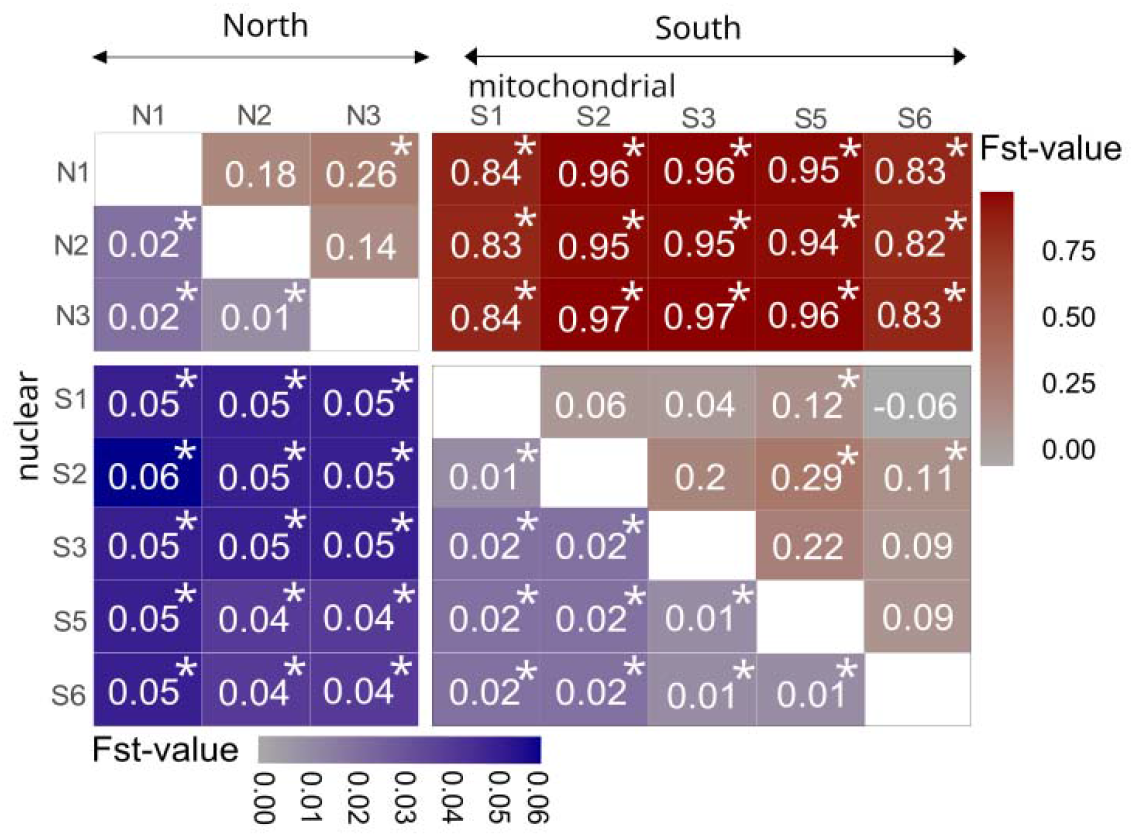
Levels of genetic differentiation between the sampled giant root-rat localities from the north and south of Bale Mountains National Park. Mitochondrial (red, above diagonal) and nuclear (blue, below diagonal) F_ST_-estimates. * in cells indicates significant differences (*p*-values < 0.05), derived by permuting haplotypes between localities for mitochondrial data and by applying a one-sample t-test on F_ST_ -values for the nuclear data.

For the nuclear data, pairwise F_ST_ - estimates between localities from different regions ranged from 0.04 to 0.06 and were higher than values between localities within regions, which ranged from 0.01 to 0.02 (Figure 4).

#### Diversity

We estimated levels of diversity for each region, and for each locality (omitting S4 as n=1). For the mitogenomes, diversity in the south (π = 0.003 ± 0.0002) was significantly higher than in the north (π = 0.001 ± 0.001, p < 0.05; Figure 2B), which reflected the presence of the divergent mitochondrial lineages in individuals WM07 and GG01 from localities S1 and S6 (Figure 2A). Thus, this was also apparent in localities S1 and S6 having the highest diversity levels, with S6 showing significantly differentiated levels of diversity (p < 0.05) and S1 showing marginal significant differentiation (p > 0.05 < 0.1, supporting information Table S3) to the remaining localities in the south.

Based on the nuclear data, we also observed significantly higher diversity in the south (π = 0.257 ± 0.0364) than in the north (π = 0.249 ± 0.0357, p < 0.05; Figure 2B). Among localities in the south, the two central localities S1 and S2 had lower diversity levels than the localities sampled further to the southeast (S3, S5, S6). All localities had significantly differentiated levels of diversity (p < 0.05, supporting information Table S3).

### 3.2 Landscape genetics

We used a reciprocal causal modelling approach with partial Mantel tests to relate genetic differentiation to the environmental distance matrices of the selected variables (vegetation index, soil moisture index, slope and elevation). Analysing levels of differentiation estimated for each region we found that elevation was the strongest model. Elevation was significantly related to genetic differentiation in all partial Mantel tests, regardless of the second environmental distance matrix controlling elevation (supporting information Table S4 A). Further, elevation showed the strongest relative support; all relative correlation values were positive, after the effect of the other environmental models were removed (supporting information Table S4 B). Geographic distance showed a significant relation with genetic differentiation when controlled by vegetation index, soil moisture index or slope, but was non-significant and the relative correlation value was negative, when controlled by elevation. We could not find significant effects of vegetation, soil moisture or slope on genetic differentiation; all variables had non-significant Mantel correlation values, showing the least support in the reciprocal causal modelling matrix (supporting information Table S4).

The raster layer optimization approach revealed that slope and elevation were the best-fitting models for explaining genetic differentiation. Slope was selected as top model in 71%, and elevation in 29% of the times across 10,000 bootstrap iterations (Table 1). The average weight from the bootstrap analyses supported geographic distance as a driver for genetic differentiation, while the parameters rank, AIC, maximum likelihood and R^2^ suggested slope and elevation as the best models (Table 1). The the response curve of the optimised resistance surface showed that the resistance costs increased with increasing slope (supporting information Figure S3 C). As in the partial Mantel tests, we could not identify a contribution of the vegetation index or soil moisture index to genetic differentiation (Table 1).

## 4. Discussion

Using complete mitochondrial genomes and low-coverage nuclear genomes from 77 giant root-rat individuals, we uncovered a clear subdivision of localities in the north and the south of the species range in the Bale Mountains, Ethiopia. Landscape genetic analysis identified topographic barriers such as the steep slopes and elevation differences between the two regions as the main drivers of population subdivision. Within regions, we did not identify any clear spatial structuring, suggesting a high level of gene flow when topographic barriers are absent.

### Genetic subdivision between regions

The significant north and south geographic subdivision in the mitochondrial and nuclear genomes, was evidenced, for instance, by a high number of substitutions separating the divergent mitochondrial lineages present in each region, or by the nuclear PCA analysis (Figure 2, supporting information Figure S1). However, two giant root-rat individuals sampled in the south were mitochondrially more closely related to their northern counterparts than their source region, which may indicate ancient gene flow. Despite their closer relationship, both had a large number of substitutions distinguishing them from the rest of the northern individuals, suggesting that ancestral mitochondrial lineages may be retained in those two individuals that are also present in the north - a remnant of their shared evolutionary history. The phylogeny indicated these two southern individuals as basal to each of two distinct haplogroups found in the north, suggesting the lineages were derived from two distinct divergence events. Ancient lineage retention in the south was supported by our nuclear analysis, which found no evidence of recent or past gene flow between north and south (Figures 2D, 3).

The presence of genetically distinct subpopulations may be attributed to long-lasting extrinsic barriers, which prevent genetic exchange between them (Avise, 2000; Bryja et al., 2010). Slope and elevation were identified as the primary drivers of genetic differentiation in our landscape analysis, and in combination they presumably cause the genetic subdivision between regions (Table 1, supporting information Table S4), similar to what has been observed in other fossorial rodents such as the Brazilian tuco tuco of the dunes (*Ctenomys flamarioni*), or the common water vole (*Arvicola terrestris*) (Berthier et al., 2005; Fernández-Stolz et al., 2007). The south comprises the Sanetti Plateau, which is ∼300 – 500 m higher in altitude than the northern region (Figure 1C) and the plateau margins northwards are characterised by broad valleys with steep slopes covered by dense *Erica* thickets. In addition to the slopes which themselves act as a barrier, the *Erica* thickets may further limit the dispersal of the species (Miehe & Miehe, 1994; Yalden, 1985). Giant root-rats are adapted to open grasslands with low vegetation and avoid dense shrubs such as *Erica*, likely due to the difficulties of burrowing in woody ground and the absence of food-plants. Additionally, the plateau is bounded by congealed lava flows of unknown age to the northwest. These barriers to dispersal and the fossorial lifestyle of the giant root-rat limiting the species’ ability to traverse pronounced topographic structures, presumably caused the strong genetic subdivision of the species. Our landscape genetic result is in agreement with recent satellite-based mapping of the giant-root rat’s distribution, which found that the texture of the landscape is the most critical factor in explaining the species’ range (Wraase et al., 2022).

In addition to slope and elevation, the pronounced subdivision observed in giant root-rats may also have been reinforced by glacial extents during the Late Pleistocene. The Bale Mountains are currently ice-free, but the Sanetti Plateau, the south region, was glaciated between ∼42,000 to 16,000 years ago (kya) (Figure 1C; Groos et al., 2021; Ossendorf et al., 2019). Except for this last glacial extent, exposure ages of moraines in the valleys in the northwestern part of the plateau (up to ∼100 kya), and stone stripes close the mountain Tullu Dimtu (up to ∼360 kya) could be interpreted in favour of earlier glacial periods (Groos et al., 2021). Possibly, giant root-rat individuals in the south were pushed towards the outer margins of the plateau by the glaciers, which increased the separation to the individuals in the north. As the glaciers retreated, colonisation of the central plateau from a more southern Late Pleistocene refugium may explain the significantly lower diversity in the central localities (S1 and S2) in comparison to the south eastern ones (S3, S5, S6, supporting information Table S3). This would be in agreement with the often-proposed hypothesis that populations of mammals exhibit reduced genetic diversity on recently deglaciated land (e.g. Hewitt, 1996, 2004). The glacial extent in the north and northwestern valleys of the plateau margins persisted until ∼16,000 years ago, while the ice shield on the plateau around Tullu Dimtu was smaller in extent already ∼20,000 years ago (Groos et al., 2021).

Although vegetation and soil moisture were previously identified as essential factors influencing the local abundance of giant root-rats (Asefa et al., 2022; Šklíba et al., 2017), our study did not indicate any effect on genetic differentiation (Table 1), suggesting these factors play less of a role in hindering gene flow at the range-wide scale. However, the spatially coarse vegetation and soil moisture indices used in our analysis may not fully capture the highly specific food and soil requirements of the giant root-rat. Their primary food resource is *Alchemilla* (Yaba et al., 2011). The vegetation index, which is based on remotely-derived satellite data, may not distinguish its spectral signal from other non-preferred plants (Wraase et al., 2022). Additionally, the giant root-rat requires soil layers of approximately 50 cm in depth to engineer burrow systems and for thermoregulation (Sillero-Zubiri et al., 1995; Šumbera et al., 2020), and while soil depth and moisture are likely correlated (deeper soil can store more water, Tromp-van Meerveld & McDonnell, 2006), soil moisture as a proxy for soil availability may not capture areas of sufficient soil depth. The vegetation and soil moisture indices were derived from a Sentinel-2 scene captured in December, just after the rainy season. During this period, vegetation is still lush and green and the soil is moist across large parts in the Bale Mountains National Park, and this thus might not fully reveal the specific habitat requirements (related to its preference for moorlands and wet grasslands with good soil depth) of the giant root-rat at that time of the year.

### Gene flow within regions

We observed a lack of structuring among localities within both regions. Levels of differentiation were low, with nuclear F_ST_ - estimates of 0.01-0.02 within regions, which is considered as weak differentiation for nuclear data (Figure 4; Weir & Cockerham, 1984; Wright, 1978). This indicates high level of dispersal and gene flow across distances of at least 16 km, which was the maximum distance between two sampling localities within regions. The ability of giant root-rats to disperse across such relatively large distances was in contrast to our expectations; giant root-rats are fossorial, solitary and territorial. We had expected this, in combination with the heterogeneity in soil structure and food availability across its range, would lead to stronger genetic structuring at small spatial scales, similar to what has been observed in other fossorial rodents (Mapelli et al., 2012; Schweizer et al., 2007). Although direct observations for this are still lacking, the limited substructuring within regions and the large dispersal distances suggest that giant root-rats can disperse aboveground and for relatively large distances. In fact, giant root-rats show morphological adaptations to surface activity, in that their eyes are situated dorsally on the head, which allows them to detect predators in open habitats (Yalden, 1985). In support of our findings, radio tracking has evidenced the dispersal of a giant root-rat individual over a distance of up to 270 m within a span of two days; the tracked individual traversed across damp soil, suggesting it did not disperse underground (Šklíba et al., 2020). Aboveground dispersal has also been documented in other fossorial, solitary rodent species, such as blind mole-rats (*Spalax microphthalmus*; Zagorodniuk et al., 2018) and Tibetian plateau zokors (*Eospalax fontanieri*; Chu et al., 2021). Even in strictly subterranean African mole-rats, long-distance dispersal is not precluded (*Fukomys damarensis*, Bathyergidae; Finn et al., 2022). For giant root-rats, aboveground dispersal attempts could be triggered by decreasing food supply, the absence of sexual partners, or the presence of competitors (Šklíba et al., 2020; Zagorodniuk et al., 2018). Also, the behaviour may circumvent the patchy availability of suitable habitats and small home-ranges, maintaining gene flow and limiting genetic structuring across small spatial scales.

Dispersal events in the giant root-rat may be male-dominated, as it has been observed in tuco tucos (*Ctenomys talarum* and *C. australis*; Cutrera et al., 2005; Mora et al., 2010), Chinese zokor (*Eospalax fontanierii*, Zhang, 2007), giant mole-rats (*F. mechowii*; Kawalika & Burda, 2007), and arvicoline rodents (Le Galliard et al., 2012). While sex-specific dispersal has not been studied in the giant root-rat yet, the observation of males being more frequently involved in dispersal attempts compared to females (Šklíba et al., 2020) and that microsatellite analysis also indicate that males disperse for longer distances (Dovičicová et al. in prep.) suggests that this type of dispersal may be prevalent in the species.

Our nuclear data did suggest slight subdivision in the south, with localities in the central part of the plateau (S1, S2) being more differentiated from localities in the southeast (S3-S6; Figure 1C). This was evidenced by increased F_ST_ in their pairwise comparisons and their segregation on the second principal component on the PCA (Figure 2C, D). Although overall differentiation among localities in the south were low, this pattern may reflect topographic features; the mountain Tullu Dimtu, the highest peak in the Bale Mountains National Park with 4,377 m a.s.l., is located close to localities S3 and S4, and may hinder gene flow (Figure 1C).

### Conservation implications

Through landscape genetic analysis, we identified the drivers of population subdivision between north and south to be topographic barriers in the form of slope and elevation. While the species is capable of dispersing locally, our findings suggest that giant root-rats in the north and south must be considered separately when developing conservation strategies, as there is no opportunity for dispersal and gene flow between them. Giant root-rat impact their surrounding environment as ecosystem engineers and primary prey for the endangered Ethiopian wolf, which underscores the importance of their persistence (Sillero-Zubiri & Gottelli, 1995; Šklíba et al., 2017). Already, the giant root-rat is believed to have a small census size due to its limited distribution range (although no census estimate is available), and is listed as endangered by the IUCN (Lavrenchenko & Kennerley, 2016). The potential for reduction of the species’ distribution range due to increasing human activities in the form of expanding livestock grazing and human settlements in the Bale Mountains, could harm the species’ persistence with negative consequences for the overall ecological balance in the region (Gashaw, 2015; Mekonen, 2020; Stephens et al., 2001). Our study yields some key insights for planning future conservation strategies for the species and highlights the value of genomic data in expanding our understanding of the population dynamics and environmental features that drive the structuring of range-limited fossorial species. With ongoing environmental changes, it is crucial to utilize this knowledge to safeguard mountain biodiversity and ecosystem functioning.

## Acknowledgements

This work was supported by the German Research Council (DFG) in the framework of the joint Ethio-European DFG Research Unit 2358 “The Mountain Exile Hypothesis. How humans benefited from and re-shaped African high-altitude ecosystems during Quaternary climate changes” [FA-925/14-1], [OP-219/10-2] and [SCHA-2085/3-1]. We are grateful to the Ethiopian Wildlife Conservation Authority, the College of Natural and Computational Sciences (Addis Ababa University), the Department of Plant Biology and Biodiversity Management (Addis Ababa University), the Frankfurt Zoological Society, the Ethiopian Wolf Project, and the Bale Mountains National Park for their cooperation and kind permission to conduct fieldwork. We are thankful to Awol Assefa, Wege Abebe, Mohammed Ahmed Muhammed and Katinka Thielsen for contributing to the preparation and implementation of the fieldwork, Christian Lampei for input on landscape genetic analyses, Alexander Groos for the input on Late Pleistocene glaciation, and Usman Abdella, Hamza Ahmed, Mohammed Kadir, Kasim Adem, Hussein Umer and Sophie Haje, for their great assistance in the field. The research was also supported by Villum Fonden Young Investigator Programme, grant no 13151 and Independent Research Fund Denmark, Sapere Aude: DFF-Forskningsleder, grant no 9064-00025B to EDL.

## Data accessibility and benefit-sharing statement

Data accessibility: The mitochondrial genome data that support the findings of this study will be openly available in GenBank of NCBI at https://www.ncbi.nlm.nih.gov under accession no. OQ207545-OQ207620 and MW751806, upon acceptance of this paper. The raw sequencing reads will be available in the associated BioProject PRJNA940645.

Benefit sharing: This project was designed as a joint Ethio-European research collaboration and developed with scientists from Ethiopia that provided the genetic samples. Our project is committed to international scientific partnerships and our collaborators are included as co-authors. The research addresses a priority concern, in this case the conservation of the studied organism.

## Author contributions

NF, GM, LO, TW and DGS designed the research concept and NF, LO and DGS secured the project funding. VMR, AA, LW and DGS captured specimens in the field. VMR and AR-I conducted lab work. VMR, AR-I and MVW analysed the data with contributions from RS, EDL, and DGS for genetic and ecological interpretation. LW generated raster layers for landscape genetic analyses. VMR, EDL and DGS wrote the manuscript with contributions from all co-authors. All authors gave final approval for publication and agreed to be held accountable for the work carried out within this publication.

