## supporting information for "Topographic barriers drive the pronounced genetic subdivision of a range-limited fossorial rodent"

**Supporting figures**

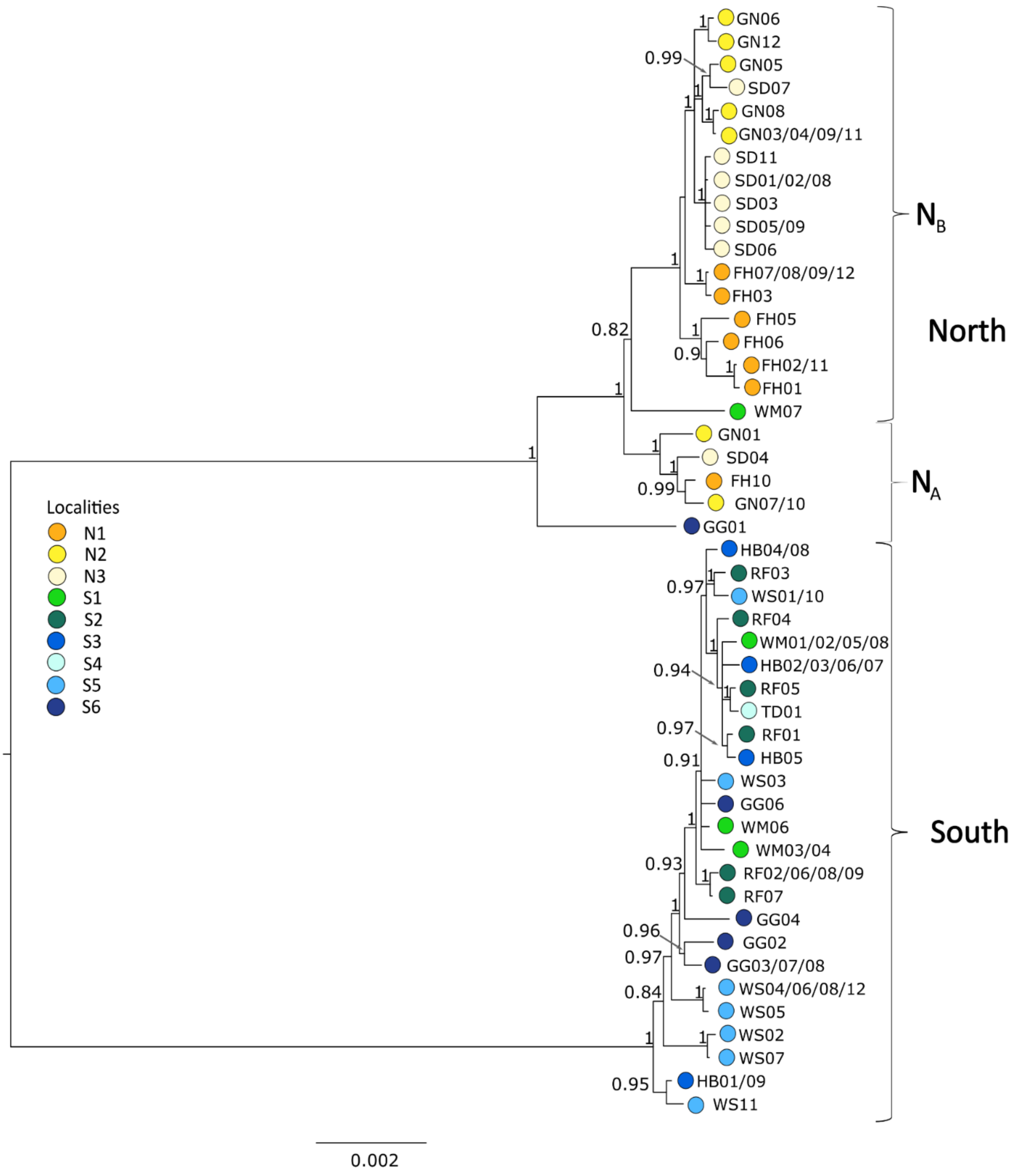


Figure S1: MrBayes phylogenetic tree of 48 unique mitochondrial haplotypes of the giant root-rat. Colours of dots show source sampling localities of individuals, see map Figure 1A; more than one sample identifier indicates shared haplotype sequences among individuals. Genetic groups discussed in the main text are indicated. Bootstrap support values > 0.8 are shown at nodes. Scale bar shows expected substitution per site.

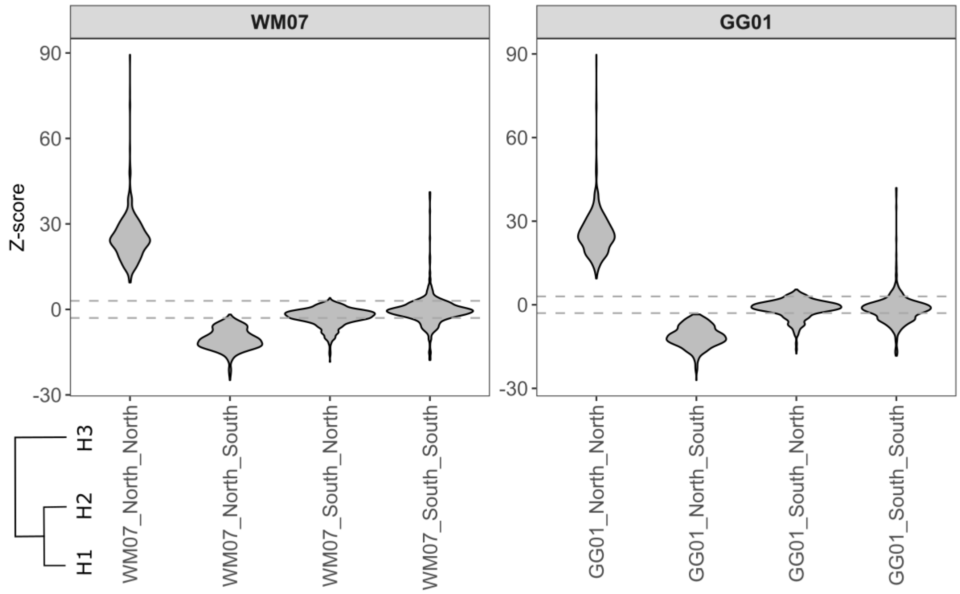


Figure S2: Z-scores of D-statistics test of gene flow analyses. Z-scores show the significance of D-scores (Figure 2, main text) from 0. |Z|>3 (shown by the dotted lines) represents a significant difference from 0, determined by a one-sample Wilcoxon signed rank test

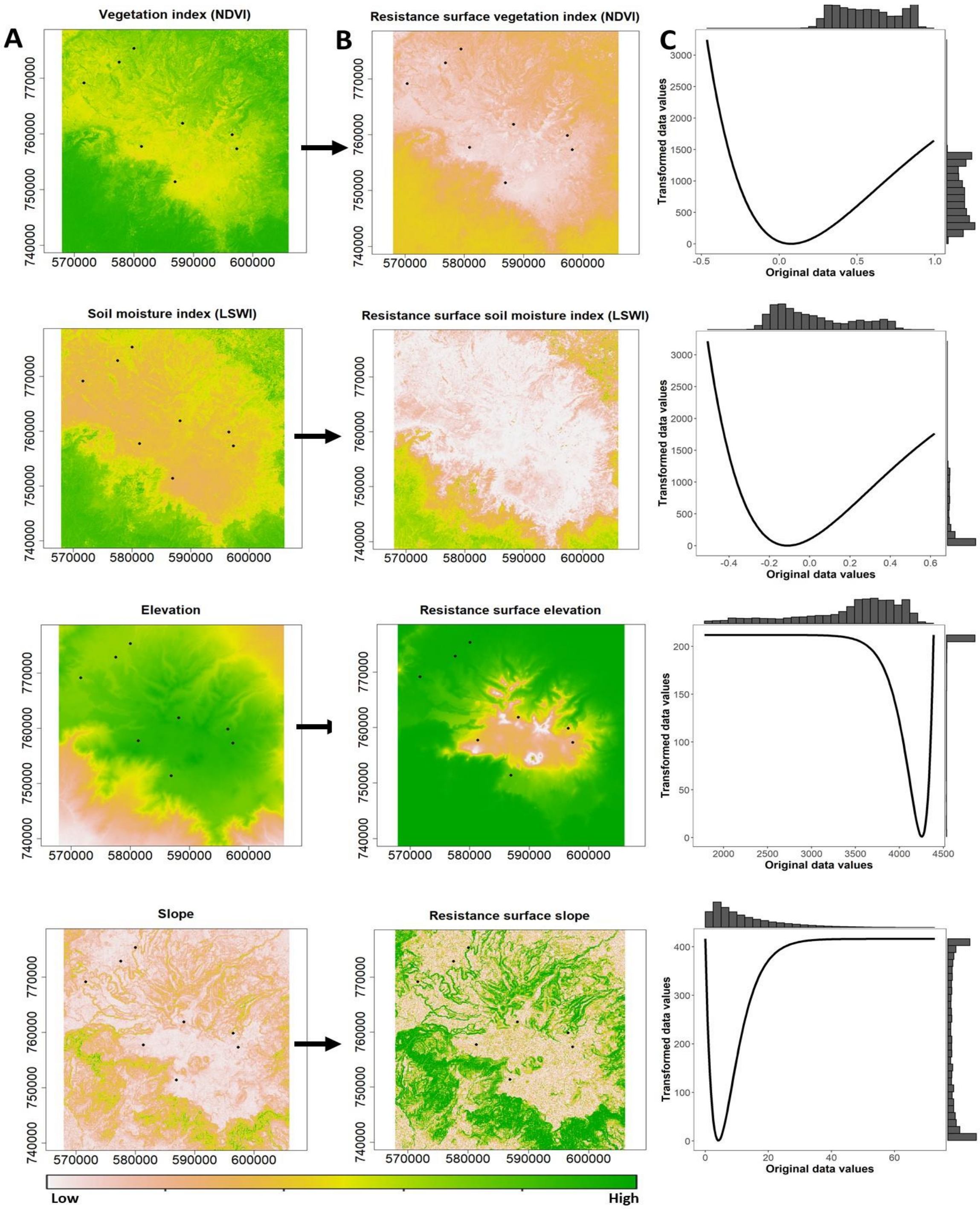


Figure S3: Raster layers of environmental variables and resistance surfaces after raster layer optimization. A) Raster layers of the Normalised Vegetation Differentiation Index (NDVI, referred to as vegetation index in main text), Land Surface Water Index (LSWI, soil moisture index), elevation, and slope. The layers were derived from a Sentinel-2 scene (www.earthexplorer.usgs.gov) (B) Resistance surfaces after the single surface optimization with ResistanceGA. The resistance costs are indicated by the colour of the bar. C) Response curves for the environmental variables, showing the resistance cost values obtained after the optimization procedure. Histogram bars indicate the frequency of each resistance value of the environmental variable.

### Supporting tables

Table S1: Sample information of the 77 giant root-rat individuals analysed, including NCBI GenBank accession number, sampling region and locality, elevation and putative sex as observed during sampling in the field.

| GenBank Accession number | Sample ID | Area | Locality | Longitude | Latitude | Elevation [m] | Sex |
|---|---|---|---|---|---|---|---|
| OQ207553 | WM01 | South | C1 | 39.7993786 | 6.89272543 | 4067 | male |
| OQ207554 | WM02 | South | C1 | 39.7979935 | 6.89250161 | 4052 | female |
| OQ207555 | WM03 | South | C1 | 39.7984543 | 6.89203954 | 4066 | female |
| OQ207556 | WM04 | South | C1 | 39.7984361 | 6.8919853 | 4074 | male |
| OQ207557 | WM05 | South | C1 | 39.7984264 | 6.89159638 | 4077 | male |
| OQ207558 | WM06 | South | C1 | 39.7982458 | 6.89185899 | 4069 | female |
| OQ207559 | WM07 | South | C1 | 39.797776 | 6.8923663 | 4061 | female |
| OQ207560 | WM08 | South | C1 | 39.7973869 | 6.89243931 | 4059 | male |
| OQ207561 | RF01 | South | S2 | 39.7341839 | 6.85683888 | 4060 | female |
| OQ207545 | RF02 | South | S2 | 39.7361359 | 6.85497257 | 4077 | female |
| OQ207546 | RF03 | South | S2 | 39.7353867 | 6.85627624 | 4068 | female |
| OQ207547 | RF04 | South | S2 | 39.7340746 | 6.85637774 | 4045 | male |
| OQ207548 | RF05 | South | S2 | 39.7359542 | 6.85453867 | 4079 | male |
| OQ207549 | RF06 | South | S2 | 39.7360538 | 6.85455661 | 4082 | female |
| OQ207550 | RF07 | South | S2 | 39.7360626 | 6.85434855 | 4085 | female |
| OQ207551 | RF08 | South | S2 | 39.7357906 | 6.85404143 | 4086 | female |
| OQ207552 | RF09 | South | S2 | 39.7359544 | 6.85463817 | 4074 | male |
| OQ207565 | FH01 | North | N1 | 39.722694 | 7.01545104 | 3432 | female |
| OQ207566 | FH02 | North | N1 | 39.723002 | 7.01585001 | 3450 | male |
| OQ207567 | FH03 | North | N1 | 39.723049 | 7.01484603 | 3454 | male |
| OQ207568 | FH05 | North | N1 | 39.724233 | 7.01355002 | 3460 | male |
| OQ207569 | FH06 | North | N1 | 39.724517 | 7.01387499 | 3461 | female |
| OQ207570 | FH07 | North | N1 | 39.724613 | 7.01381296 | 3463 | female |
| OQ207571 | FH08 | North | N1 | 39.724612 | 7.01382796 | 3466 | male |

| | | | | | | | |
|---|---|---|---|---|---|---|---|
| OQ207572 | FH09 | North | N1 | 39.724365 | 7.01379502 | 3462 | NA |
| OQ207562 | FH10 | North | N1 | 39.724167 | 7.01372101 | 3460 | NA |
| OQ207563 | FH11 | North | N1 | 39.72443 | 7.01378597 | 3461 | male |
| OQ207564 | FH12 | North | N1 | 39.72443 | 7.01378597 | 3461 | NA |
| OQ207573 | GG01 | South | S6 | 39.873655 | 6.87388103 | 3996 | female |
| OQ207574 | GG02 | South | S6 | 39.873954 | 6.87302599 | 3932 | female |
| OQ207575 | GG03 | South | S6 | 39.873791 | 6.87366704 | 4039 | male |
| OQ207576 | GG04 | South | S6 | 39.873474 | 6.87347903 | 4045 | female |
| MV751806 | GG06 | South | S6 | 39.873731 | 6.87346403 | 4039 | female |
| OQ207577 | GG07 | South | S6 | 39.873964 | 6.873229 | 4039 | female |
| OQ207578 | GG08 | South | S6 | 39.874289 | 6.87294401 | 4042 | female |
| OQ207587 | GN01 | North | N2 | 39.649925 | 6.95700897 | 3330 | male |
| OQ207588 | GN03 | North | N2 | 39.649347 | 6.95741398 | 3543 | male |
| OQ207589 | GN04 | North | N2 | 39.648648 | 6.95798303 | 3546 | female |
| OQ207579 | GN05 | North | N2 | 39.650556 | 6.958222 | 3446 | male |
| OQ207580 | GN06 | North | N2 | 39.648671 | 6.95795302 | 3544 | female |
| OQ207581 | GN07 | North | N2 | 39.651052 | 6.95835804 | 3546 | female |
| OQ207582 | GN08 | North | N2 | 39.650912 | 6.95715699 | 3550 | female |
| OQ207583 | GN09 | North | N2 | 39.650212 | 6.95687796 | 3550 | male |
| OQ207584 | GN10 | North | N2 | 39.650925 | 6.95836902 | 3550 | male |
| OQ207585 | GN11 | North | N2 | 39.648578 | 6.95802502 | 3548 | female |
| OQ207586 | GN12 | North | N2 | 39.651665 | 6.95825301 | 3552 | male |
| OQ207590 | HB01 | South | S2 | 39.7853 | 6.79550101 | 3623 | female |
| OQ207591 | HB02 | South | S2 | 39.78964 | 6.79768098 | 3853 | female |
| OQ207592 | HB03 | South | S2 | 39.784437 | 6.79520002 | 3850 | male |
| OQ207593 | HB04 | South | S2 | 39.787152 | 6.79719399 | 3852 | female |
| OQ207594 | HB05 | South | S2 | 39.789308 | 6.797743 | 3851 | female |
| OQ207595 | HB06 | South | S2 | 39.78687 | 6.79702501 | 3850 | female |
| OQ207596 | HB07 | South | S2 | 39.787018 | 6.79715199 | 3854 | male |
| OQ207597 | HB08 | South | S2 | 39.783954 | 6.79434397 | 3849 | female |
| OQ207598 | HB09 | South | S2 | 39.784998 | 6.79543303 | 3849 | male |
| OQ207600 | SD01 | North | N3 | 39.702559 | 6.99164601 | 3509 | male |
| OQ207601 | SD02 | North | N3 | 39.702437 | 6.99169002 | 3477 | NA |
| OQ207602 | SD03 | North | N3 | 39.702509 | 6.99163897 | 3508 | NA |

| | | | | | | | |
|---|---|---|---|---|---|---|---|
| OQ207603 | SD04 | North | N3 | 39.70244 | 6.99198204 | 3501 | female |
| OQ207604 | SD05 | North | N3 | 39.701275 | 6.991054 | 3394 | female |
| OQ207605 | SD06 | North | N3 | 39.702295 | 6.99160896 | 3507 | male |
| OQ207606 | SD07 | North | N3 | 39.702515 | 6.99174198 | 3516 | NA |
| OQ207607 | SD08 | North | N3 | 39.70168 | 6.99056902 | 3506 | female |
| OQ207608 | SD09 | North | N3 | 39.701255 | 6.991054 | 3502 | female |
| OQ207599 | SD11 | North | N3 | 39.701205 | 6.98984399 | 3503 | male |
| OQ207609 | TD1 | South | S4 | 39.826091 | 6.82700104 | 4224 | male |
| OQ207610 | WS01 | South | S5 | 39.88116 | 6.850291 | 4117 | male |
| OQ207611 | WS02 | South | S5 | 39.881139 | 6.85024498 | 4114 | female |
| OQ207612 | WS03 | South | S5 | 39.881217 | 6.850794 | 4100 | female |
| OQ207613 | WS04 | South | S5 | 39.880863 | 6.850406 | 4111 | female |
| OQ207614 | WS05 | South | S5 | 39.880683 | 6.85055603 | 4111 | female |
| OQ207615 | WS06 | South | S5 | 39.880764 | 6.85039301 | 4114 | male |
| OQ207616 | WS07 | South | S5 | 39.881243 | 6.85041497 | 4113 | female |
| OQ207617 | WS08 | South | S5 | 39.88116 | 6.85078603 | 4111 | NA |
| OQ207618 | WS10 | South | S5 | 39.881335 | 6.85093397 | 4112 | NA |
| OQ207619 | WS11 | South | S5 | 39.880807 | 6.850924 | 4113 | male |
| OQ207620 | WS12 | South | S5 | 39.881034 | 6.84992597 | 4116 | male |

Table S2: Mapping statistics of the 77 giant root-rat samples analysed. Raw sequencing reads were mapped against the hoary bamboo rat (*Rhizomys pruinosus*) nuclear genome (Genbank accession: VZQC00000000.1; Guo et al. 2021) combined with the giant root-rat mitogenome (Genbank accession: MW751806; Reuber et al. 2021). Bp = base pairs; Mitogenomes = mitochondrial genomes

| **Sample ID** | **# Raw read pairs** | **# Reads mapping** | **Nuclear coverage** | **Nuclear mapped bp** | **Mitogenome coverage** | **Mitogenome mapped bp** |
|---|---|---|---|---|---|---|
| FH01 | 16,978,992 | 7,992,025 | 0.29 | 1,080,644,275 | 226.90 | 3,776,908 |
| FH10 | 18,494,391 | 12,855,168 | 0.51 | 1,901,006,013 | 182.64 | 3,040,203 |
| FH11 | 19,750,607 | 10,262,289 | 0.40 | 1,487,884,552 | 144.36 | 2,403,072 |
| FH12 | 17,227,253 | 11,333,662 | 0.45 | 1,651,542,774 | 128.76 | 2,143,278 |
| FH02 | 14,883,520 | 8,146,465 | 0.32 | 1,183,176,147 | 168.19 | 2,799,747 |
| FH03 | 16,251,258 | 10,476,801 | 0.43 | 1,582,232,753 | 181.38 | 3,019,192 |
| FH05 | 16,852,614 | 10,499,672 | 0.41 | 1,510,446,294 | 250.02 | 4,161,839 |
| FH06 | 19,896,506 | 12,832,141 | 0.52 | 1,906,513,326 | 192.50 | 3,204,294 |
| FH07 | 19,431,703 | 9,452,783 | 0.35 | 1,283,194,137 | 147.39 | 2,453,482 |
| FH08 | 15,914,201 | 8,510,001 | 0.32 | 1,189,432,482 | 179.62 | 2,990,019 |
| FH09 | 17,438,587 | 9,371,585 | 0.36 | 1,320,089,804 | 234.83 | 3,908,968 |
| GG01 | 22,295,022 | 13,577,661 | 0.55 | 2,023,849,334 | 170.93 | 2,845,211 |
| GG02 | 17,747,863 | 11,139,869 | 0.44 | 1,630,575,444 | 147.06 | 2,447,920 |
| GG03 | 16,768,929 | 7,684,884 | 0.27 | 1,014,360,345 | 102.97 | 1,714,101 |
| GG04 | 24,563,197 | 1,535,631 | 0.12 | 442,335,192 | 24.94 | 415,083 |
| GG06 | 24,526,276 | 12,539,983 | 0.49 | 1,803,094,277 | 220.69 | 3,673,638 |
| GG07 | 20,381,975 | 12,354,185 | 0.49 | 1,793,309,550 | 232.45 | 3,869,412 |
| GG08 | 4,425,384 | 12,274,902 | 0.47 | 1,719,823,251 | 212.03 | 3,529,394 |
| GN05 | 26,875,116 | 7,969,437 | 0.38 | 1,396,716,389 | 94.19 | 1,567,817 |
| GN06 | 19,415,999 | 10,070,286 | 0.40 | 1,474,915,866 | 155.88 | 2,594,709 |
| GN07 | 16,723,334 | 7,006,642 | 0.28 | 1,042,333,258 | 121.03 | 2,014,703 |
| GN08 | 25,301,937 | 10,848,812 | 0.43 | 1,602,419,125 | 191.59 | 3,189,169 |
| GN09 | 21,256,957 | 9,479,533 | 0.38 | 1,401,602,300 | 155.62 | 2,590,518 |
| GN01 | 16,003,381 | 8,207,272 | 0.32 | 1,175,154,307 | 230.38 | 3,834,928 |
| GN10 | 21,011,631 | 13,404,421 | 0.53 | 1,943,213,143 | 218.10 | 3,630,495 |
| GN11 | 25,847,517 | 16,396,946 | 0.64 | 2,378,222,263 | 242.29 | 4,033,095 |
| GN12 | 22,475,014 | 12,581,467 | 0.50 | 1,831,406,864 | 146.60 | 2,440,378 |
| GN03 | 14,555,865 | 6,320,660 | 0.24 | 895,094,428 | 143.31 | 2,385,457 |
| GN04 | 19,336,769 | 12,560,208 | 0.50 | 1,835,220,325 | 240.73 | 4,007,196 |
| HB01 | 21,973,877 | 10,182,380 | 0.43 | 1,583,136,018 | 102.69 | 1,709,392 |
| HB02 | 25,529,934 | 17,319,108 | 0.68 | 2,504,372,366 | 219.52 | 3,654,153 |
| HB03 | 21,702,035 | 7,539,986 | 0.30 | 1,122,205,194 | 129.43 | 2,154,491 |

| | | | | | | |
|---|---|---|---|---|---|---|
| HB04 | 23,433,747 | 14,126,631 | 0.56 | 2,088,257,975 | 159.68 | 2,658,019 |
| HB05 | 23,994,437 | 14,449,681 | 0.57 | 2,118,929,629 | 361.16 | 6,011,848 |
| HB06 | 19,951,156 | 11,088,337 | 0.44 | 1,626,922,114 | 126.54 | 2,106,308 |
| HB07 | 16,010,826 | 9,260,591 | 0.37 | 1,356,991,234 | 160.97 | 2,679,465 |
| HB08 | 24,162,995 | 13,835,736 | 0.56 | 2,054,483,977 | 182.77 | 3,042,383 |
| HB09 | 23,621,873 | 13,184,020 | 0.57 | 2,118,929,629 | 199.79 | 3,325,660 |
| SD01 | 21,121,011 | 6,635,909 | 0.30 | 1,097,745,936 | 144.43 | 2,404,109 |
| SD11 | 14,776,896 | 6,401,618 | 0.26 | 952,603,757 | 85.26 | 1,419,154 |
| SD02 | 20,060,768 | 13,430,376 | 0.54 | 1,982,436,708 | 249.92 | 4,160,133 |
| SD03 | 12,197,516 | 8,320,803 | 0.33 | 1,211,921,153 | 183.82 | 3,059,800 |
| SD04 | 15,072,836 | 9,636,470 | 0.38 | 1,394,660,022 | 171.47 | 2,854,279 |
| SD05 | 3,300,973 | 10,258,149 | 0.40 | 1,466,709,062 | 258.30 | 4,299,641 |
| SD06 | 18,112,628 | 11,862,203 | 0.47 | 1,728,272,832 | 291.72 | 4,855,990 |
| SD07 | 17,365,379 | 4,706,885 | 0.19 | 711,542,791 | 93.47 | 1,555,961 |
| SD08 | 14,927,338 | 7,899,143 | 0.31 | 1,127,141,716 | 226.73 | 3,774,112 |
| SD09 | 14,754,546 | 3,961,884 | 0.22 | 814,631,762 | 116.97 | 1,947,051 |
| TD01 | 18,360,169 | 8,194,229 | 0.29 | 1,076,743,466 | 182.60 | 3,039,521 |
| WS01 | 27,711,489 | 6,982,153 | 0.43 | 1,576,780,562 | 383.22 | 6,379,103 |
| WS10 | 21,495,541 | 10,268,532 | 0.38 | 1,405,339,675 | 276.23 | 4,598,081 |
| WS11 | 20,698,366 | 12,000,838 | 0.47 | 1,726,480,366 | 153.60 | 2,556,859 |
| WS12 | 20,877,111 | 9,511,682 | 0.34 | 1,240,687,498 | 167.02 | 2,780,161 |
| WS02 | 20,283,118 | 10,724,014 | 0.42 | 1,558,731,684 | 129.18 | 2,150,311 |
| WS03 | 22,665,497 | 12,505,010 | 0.49 | 1,814,919,219 | 157.34 | 2,619,039 |
| WS04 | 20,531,719 | 11,151,303 | 0.44 | 1,626,149,287 | 148.33 | 2,469,173 |
| WS05 | 19,546,437 | 10,341,864 | 0.40 | 1,495,657,674 | 147.86 | 2,461,306 |
| WS06 | 22,699,761 | 13,505,918 | 0.54 | 1,983,859,527 | 231.53 | 3,854,088 |
| WS07 | 19,914,784 | 12,277,532 | 0.48 | 1,789,796,875 | 123.11 | 2,049,223 |
| WS08 | 17,942,128 | 10,381,806 | 0.42 | 1,552,684,778 | 126.66 | 2,108,292 |
| WM01 | 25,415,257 | 19,240,329 | 0.74 | 2,752,021,342 | 121.31 | 2,019,292 |
| WM02 | 22,638,733 | 15,910,096 | 0.65 | 2,414,133,391 | 125.53 | 2,089,639 |
| WM03 | 24,802,453 | 18,354,244 | 0.77 | 2,837,925,408 | 177.23 | 2,950,212 |
| WM04 | 22,795,120 | 14,958,043 | 0.70 | 2,582,763,905 | 158.99 | 2,646,611 |
| WM05 | 19,230,604 | 18,829,904 | 0.55 | 2,047,322,685 | 68.19 | 1,135,121 |
| WM06 | 21,399,645 | 17,553,235 | 0.63 | 2,345,642,536 | 70.48 | 1,173,188 |
| WM07 | 20,423,503 | 14,526,040 | 0.62 | 2,290,090,671 | 97.40 | 1,621,323 |
| WM08 | 22,856,570 | 17,206,734 | 0.69 | 2,568,908,319 | 93.53 | 1,556,848 |
| RF01 | 22,228,252 | 19,833,972 | 0.67 | 2,471,394,141 | 104.29 | 1,735,935 |
| RF02 | 24,670,699 | 17,585,365 | 0.71 | 2,626,112,333 | 108.40 | 1,804,460 |

| | | | | | | |
|---|---|---|---|---|---|---|
| RF03 | 19,537,627 | 20,573,370 | 0.59 | 2,187,672,230 | 116.29 | 1,935,731 |
| RF04 | 23,599,367 | 18,644,450 | 0.69 | 2,553,480,252 | 166.93 | 2,778,720 |
| RF05 | 17,913,386 | 14,957,434 | 0.56 | 2,063,675,254 | 128.10 | 2,132,328 |
| RF06 | 22,795,981 | 17,160,550 | 0.70 | 2,596,803,976 | 126.18 | 2,100,317 |
| RF07 | 22,941,232 | 16,563,368 | 0.65 | 2,408,176,843 | 99.55 | 1,657,191 |
| RF08 | 18,119,643 | 18,563,076 | 0.54 | 2,001,105,976 | 136.84 | 2,277,795 |
| RF09 | 20,711,128 | 17,982,889 | 0.64 | 2,372,186,376 | 139.21 | 2,317,277 |

Table S3: P-Values of the test of significant differences in nucleotide diversity difference between localities for mitochondrial and nuclear genomes. For mitochondrial genomes (above diagonal) a permutation approach was conducted, where haplotypes were resampled (n=1000) across the entire giant root-rat population, in order to test whether the observed genetic diversity of each locality was equal or greater than the genetic diversity simulated in the permutation approach. For nuclear genomes (below diagonal), we used a Welch-test (unpaired t-test), accounting for unequal variance. p-values < 0.05 present significant nucleotide differences between localities (in italics)

| | **N1** | **N2** | **N3** | **S1** | **S2** | **S3** | **S5** | **S6** |
|---|---|---|---|---|---|---|---|---|
| **N1** | -- | 0.804 | 0.725 | 0.069 | 0.672 | 0.759 | 0.979 | *0.042* |
| **N2** | *<0.001* | -- | 0.599 | 0.072 | 0.576 | 0.648 | 0.782 | 0.056 |
| **N3** | *<0.001* | *0.023* | -- | 0.064 | 0.897 | 0.987 | 0.746 | *0.032* |
| **S1** | 0.708 | *0.002* | *<0.001* | -- | 0.065 | 0.078 | 0.052 | 0.613 |
| **S2** | *0.001* | *<0.001* | *<0.001* | *0.003* | -- | 0.823 | 0.677 | *0.030* |
| **S3** | *<0.001* | *<0.001* | *<0.001* | *<0.001* | *<0.001* | -- | 0.755 | *0.042* |
| **S5** | *<0.001* | *<0.001* | *<0.001* | *<0.001* | *<0.001* | *0.003* | -- | *0.043* |
| **S6** | *<0.001* | *<0.001* | *<0.001* | *<0.001* | *<0.001* | *<0.001* | *<0.001* | -- |

Table S4: a) Correlation matrix of partial Mantel test of column variables (focal model) controlled by the row variables (alternative model) for mitochondrial data. a) Shown is the correlation coefficient r, bold values indicate a significant correlation. b) Reciprocal causal modelling matrix, showing the relative support of the focal models in columns (e.g. elevation distance|geographic distance), compared to alternative models in rows (geographic distance|elevation distance). Values depict differences in r-values between focal and alternative model. Positive values support the focal, negative the alternative model.

| | **Focal model** | | | | |
|---|---|---|---|---|---|
| **Alternative model** | Geographic distance | Vegetation distance | Moisture distance | Elevation distance | Slope distance |
| *A) Correlation matrix* | | | | | |
| Geographic distance | 0.00 | 0.11 | -0.06 | **0.42** | 0.21 |
| Vegetation distance | **0.56** | 0.00 | 0.18 | **0.65** | 0.19 |
| Moisture distance | **0.55** | 0.18 | 0.00 | **0.64** | **0.27** |
| Elevation distance | 0.19 | 0.16 | -0.09 | 0.00 | 0.17 |
| Slope distance | **0.55** | -0.02 | -0.19 | **0.63** | 0.00 |
| *B) Reciprocal causal modelling matrix* | | | | | |
| Geographic distance | 0.00 | -0.44 | -0.61 | 0.24 | -0.34 |
| Vegetation distance | 0.44 | 0.00 | 0.00 | 0.49 | 0.21 |
| Moisture distance | 0.61 | 0.00 | 0.00 | 0.73 | 0.46 |
| Elevation distance | -0.24 | -0.49 | -0.73 | 0.00 | -0.47 |
| Slope distance | 0.34 | -0.21 | -0.46 | 0.47 | 0.00 |
